# Exploiting Intrinsic Noise for Heterogeneous Cell Control Under Time Delays and Model Uncertainties

**DOI:** 10.1101/2023.10.07.561335

**Authors:** M P May, B Munsky

## Abstract

The majority of previous research in synthetic biology has focused on enabling robust control performance despite the presence of noise, while the understanding for how controllers may exploit that noise remains incomplete. Motivated by Maxwell’s Demon, we previously proposed a cellular control regime in which the exploitation of stochastic noise can break symmetry between and allow for specific control of multiple cells using a single input signal (i.e., single-input-multiple-output or SIMO control). The current work extends that analysis to include uncertain stochastic systems where system dynamics are are affected by time delays, intrinsic noises, and model uncertainty. We find that noise-exploiting controllers can remain highly effective despite coarse approximations to the model’s scale or incorrect estimations or extrinsic noise in key model parameters, and these controllers can even retain performance under substantial observer or actuator time delays. We also demonstrate how SIMO controllers could drive multi-cell systems to follow different trajectories with different phases and frequencies. Together, these findings suggest that noise-exploiting control should be possible even in the practical case where models are always approximate, where parameters are always uncertain, and where observations are corrupted by errors.

## 1. Introduction

Uncertain fluctuations, or ‘noise,’ is a common theme throughout many fields of engineering, and robust control is a frequent concern when attempting to control any human-made system such as vehicles [1], chemical processes [2], or biology [3]. Noise arrises from many sources, occasionally from quantum events and thermally-induced fluctuations, but more commonly from unknown or uncharacterized physical processes, and as such, these fluctuations usually cannot be efficiently reproduced or predicted using purely physical or mechanistic models. Rather, statistical models are usually employed where key mechanisms or dynamics are subject to stochastic variations or inputs that act as a proxy for random or un-modeled fluctuations elsewhere in the system. In this context, much effort has been placed on enhancing the robustness of controlled processes, where uncertain or unpredictable variations in internal or external parameters are modeled as intrinsic or extrinsic noise [4]. For many macroscopic, human-engineered systems, the resulting system dynamics can be modeled effectively simply by combining a deterministic (i.e., noise-free) model with additive noise (usually assumed to be Gaussian under arguments based on the central limit theorem) [5], and most current approaches seek to control the system to minimize variations that result from the noisy inputs.

Analyses of noise and control theory are equally relevant to understand the basic biological processes of gene transcription regulation [6]or mRNA translation regulation [7]or to modify these processes for practical use [8, 9]. However, at this mesoscopic scale of cellular biology, where transcription factors compete to activate or deactivate individual genes or where mRNA or protein molecules are present at just a few copies (or none at all) per cell, the additive noise model is much less realistic. In this case, Brownian motion, discrete stochastic gene regulation, and noisy mRNA dynamics collectively generate a fundamentally stochastic environment that cells must effectively manage. At this scale, the order or timing of a single reaction event (e.g., the binding of a transcription factor to a promoter) can have dramatic consequences that could last for several cellular generations (e.g., the activation or repression of a gene that promotes unfettered cell growth and differentiation). The cell’s drive towards homeostasis requires dealing with the inherently chaotic and noisy processes that reside within it, and despite these challenges, cells generally demonstrate strong capability to survive these noisy processes. When seeking to understand how such systems evolve or react, the central limit theorem and the Gaussian noise may not apply, and a more detailed statistical analysis of probabilistic behavior is needed uncover hidden properties of cellular control mechanisms.

One emerging field that is particularly dependent on the integration of control and noise is synthetic biology, which aims to develop modular[10] and orthogonal [11] components to sense and manipulate [12] complex logical systems, enabling them to exhibit a wide range of advanced biological behaviors[13]. Advances in optogenetics have enhanced the ability to reliably actuate embedded systems within cells, offering the potential to exert precise temporal and spatial control on cellular components[12, 9, 14, 15]. These developments have facilitated computer-programmable regulation of cellular protein production through external optogenetic inputs and smart microscope techniques[16, 17, 18]. These digital-synthetic actuators enable fine-tuned, computer-modulated control of cellular systems, previously unattainable, with faster response times compared to chemical diffusion[19, 9]. Classical and modern control methods like PID control and model predictive control have been implemented in such systems [20]to control synthetic systems to different stable points.

It has recently been shown that new control techniques that leverage the complete probability distribution information of the system could actually harness the noise of single-cell gene regulation to achieve more complicated control objectives. For example, inspired by the genetic toggle switch from Kobayashi et al.[21], Szymanska et al. [22] showed that noise could be exploited to achieve independent control of multiple cells using a single input, even despite uncertain parameters or time delays due to maturation of fluorescent proteins or limited observation of the regulatory proteins. In May et al. [23], we identified a simplified stochastic model to reproduce data measured in Baumschlager, et al. [9] for the expression from a transcription promoter under the optogenetic control of a UV-activated T7 polymerase (see model in Figure 1A, top promoter). We then proposed the addition of a positive auto-regulation (Figure 1A, bottom promoter) to help maintain an elevated expression phenotype in the presence of UV excitation, and we demonstrated how a Single-Input-Multiple-Output (SIMO) multicellular controller could control multiple cells to arbitrary phenotypes using only a single input.

**Figure 1.**
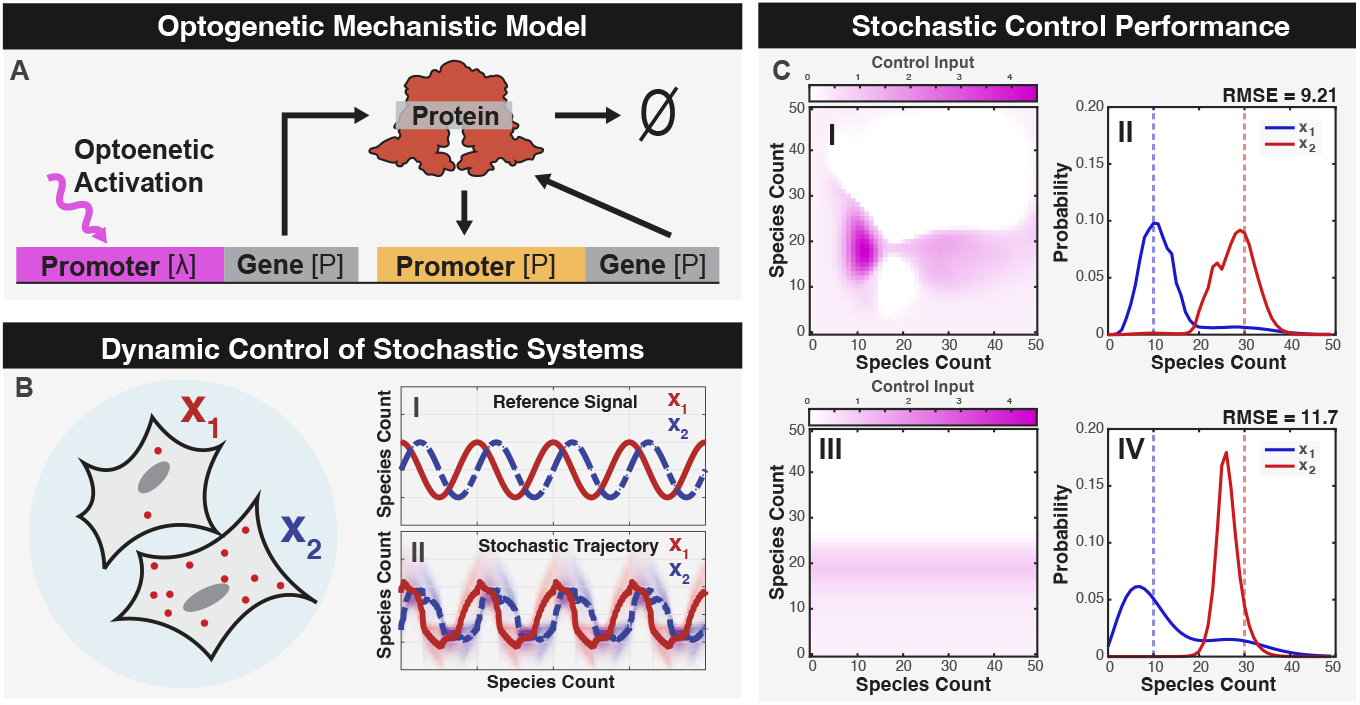
Single-input-multiple-output (SIMO) control of multiple cells using a single optogenetic input. (**A**) Schematic of the light-activated genetic system with auto-regulation. (**B**) Diagram of the stochastic SIMO control problem using two optogenetic cells sharing a single input. (**C**) Noise exploiting controllers were optimized to define a fully aware control input (I) and a partially aware control input (III), where the control signal (color scale) depends on the observed expression within the cell or cells (x- and y-axes). (II and IV) Corresponding steady state marginal distributions for the different cells (red and blue) under these controllers demonstrate a clear a break in symmetry. Dashed lines represent the control target objective. Control performance (RMSE) is show above each distribution.

This paper extends the analysis of the SIMO multicellular control problem by examining the impact of model uncertainties and fluctuating control objectives on the control performance. These uncertainties include coarse-grained approximations of the system dynamics, errors or extrinsic variations in the system parameter, and time delays between the observation of the cellular dynamics and actuation of the control process. In the following ‘Methods’ we introduce our formulation of the chemical master equation (CME) to analyze the discrete stochastic distribution of cellular responses; we define multiple controllers and demonstrate the computation of their effects on cell dynamics; and we show how the control law can be optimized to improve performance. In the ‘Results’ section, we explore the how model approximations, parameter inaccuracies, and time delays affect control performance, and we demonstrate a simple scheme for controlling cells to track a dynamically changing reference signal. Using discrete stochastic models based on the chemical master equation, we demonstrate that combining biochemical noise, nonlinear auto-regulation, and a single optogenetic feedback could control two genetically identical cells with different initial conditions to follow different desired trajectories at different frequencies and phases.

## 2. Methods

In May et al. [23], we developed two models for the description of an optogenetically controlled gene expression system. These first model consisted of six species to describe the light-activated association of two T7 split domains (species 1 and 2) which combine to form an active T7 polymerase (species 3) under optogenetic excitation. The active polymerase could then associate with inactive T7 promoters (species 4), resulting in the formation of an active allele (species 5) that could then transcribe and translated to produce the desired protein product (species 6). The second, much simpler, single-species model was developed by assuming quasi-steady equilibrium for the first five species. Both models were independently parameterized using the same experimental data from Baumschlager et al. [9]. Furthermore, an extension was made to each model to incorporate a secondary self-activated promoter-gene construct, where the expression rate was determined by a Hill function 1. Through simulations, we showed that a feedback control law could be designed to force multiple cells to different and individually chosen equilibrium states using a single optogenetic control signal. We also showed that when this control law was parametrized using the simple model, it could be used effectively to control the behavior of the more complicated system, thus demonstrating that control performance could remain high despite inaccuracies in the model. In the current work, we adopt the simpler of the two models and extend our analyses to consider the effects of additional model inaccuracies, including coarse-grained model approximations, parameter errors, and time delays, and we also explore the possibility that a single optogenetic control signal could drive the system to track temporally-changing reference signals. Section 2.1 introduces the model; Section 2.2 develops a Master Equation description of the models probabilistic dynamics; Section 2.3 defines a control objective and optimizes that metric to obtain a baseline control law; and Sections 2.4 and 2.5 introduce uncertainties into the model related to the granularity of the model approximation or the introduction of time delays, respectively.

### 2.1. Model

To assess the impact of model approximations on the implementation of noise-enhanced control strategies, we begin with the one-species model proposed by May et al. [23] (Figure. 1A). This model comprises two reactions for production and degradation of the key protein. The nonlinear, UV-dependent production is defined by the following equation:

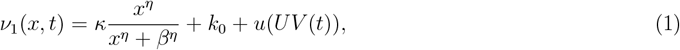

where *x* is the instantaneous protein level; *κ* is the maximum strength of the auto-regulation promoter; *β* is the concentration at which auto-regulation promoter reaches its half maximal strength; *η* is the cooperativity in the auto-regulation promoter; *k*_0_ is the leakage rate from both promoters; and *u*(*UV* (*t*)) is the UV-dependent strength of the T7 promoter. Feedback enables the external modulation of the light input using the state of the system, thereby controlling the T7 promoter strength as a function of state rather than time and eliminating the explicit time dependance in *u*(*UV* (*t*)). Protein degradation is assumed to be a first order process with rate *γ*:

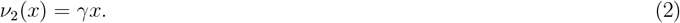

All baseline parameters describing the auto-regulation promoter, *κ, η, β*, and *k*_0_, and the degradation rate *γ* are presented in Table 1 [23] and are fixed throughout the current study.

**Table 1.** Baseline Model Parameters.

| Parameter | Value | Units | Meaning |
| --- | --- | --- | --- |
| $\kappa$ | 0.406 | 1/min | maximal production rate |
| $\beta$ | 20.0 | molecules | concentration at half-max activation |
| $\eta$ | 8.00 | unitless | cooperativity factor |
| $k_0$ | 0.0001 | 1/min | promote leakage rate |
| $\gamma$ | 0.0203 | 1/min | degradation/dilution rate |
| $u(t)$ | variable | 1/min | applied control signal |

### 2.2. Stochastic analyses of the model

To describe the discrete stochastic behavior of the above model for a population of *N*_c_ cells, we define the current state of the system as the tuple of the non-negative numbers of proteins in each cell: 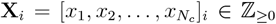, where the index *i* denotes the enumeration of the state within the countably infinite set of all possible states, i.e., **X**_*i*_ *∈ X* = *{***X**_1_, **X**_2_, …*}*. The stoichiometry vector, **s**_*μ*_, for reaction number *μ* is then defined as the change in state following that reaction event (e.g., **X**_*i*_ *→* **X**_*i*_ + **s**_*μ*_). Specifically, the 2*N*_*c*_ possible reactions are defined in pairs corresponding to production (*μ ∈ {*1, 3, 5, …*}*) and degradation (*μ ∈ {*2, 4, 6, …*}*) as:

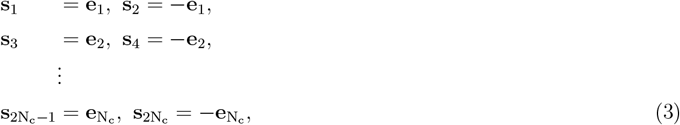

where each 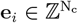 is is a Euclidean vector (i.e., unity for the *i*^th^ entry and otherwise zero). The corresponding propensity functions are:

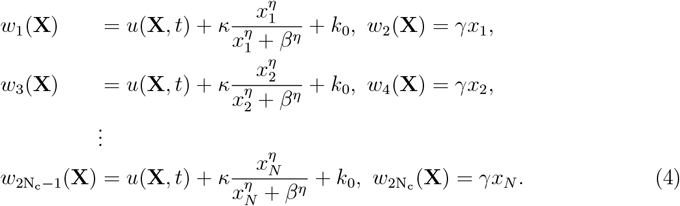

These definitions of the stoichiometry and propensity functions allow us to implement the Gillespie stochastic simulation algorithm (SSA) [24, 25] to generate representative trajectories of the stochastic process. At each step, two random numbers are generated to determine the time and the type of the next reaction. Given the current state **X**, the time until the next reaction is distributed according to an exponential random with rate parameter equal to the inverse of the sum of the propensity functions:

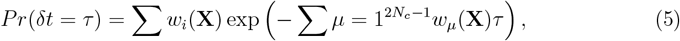

and an instance of this random variable, *δt*, can be sampled from this distribution using the expression:

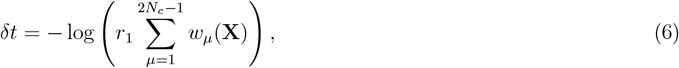

where *r*_1_ is a uniform random variable between zero and one. The probability for the specific individual reaction *R*_*k*_ to fire from all possible reactions is given by:

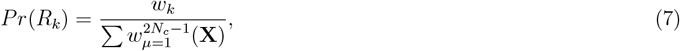

and which specific reaction that fires at time *t*+*τ* is a categorical random variable pulled from *Pr*(*R*_*k*_). The SSA is then simulated by stepping through time one reaction at a time and updating the state of the system by adding the stoichiometry of the reaction to the state of the system.

However, a more direct analysis of the chemical master equation (CME) is necessary to quantify performance and optimize the controller design. The high dimensional CME is a linear ODE that describes the time-dependent changes in probability mass of all possible states. Using the specified reaction propensities and stoichiometries, the CME can be expressed as:

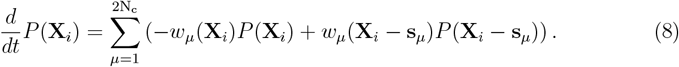

For convenience, the CME can also be formulated more compactly in matrix-vector form as:

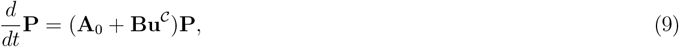

where **P** = [*P* (**X**_1_), *P* (**X**_2_), …]^*T*^ is the enumerated probability mass vector for all possible states of the system; **A**_0_ is the infinitesimal generator of the stochastic process due to the autoregulation promoter and degradation events; **u**^*𝒞*^ = [*u*^*𝒞*^(**X**_1_), *u*^*𝒞*^(**X**_2_), …]^*T*^ is the collection of control inputs associated with each state; and **Bu**^*𝒞*^ is the contribution that these control inputs make to the infinitesimal generator when included into the feedback process. More specifically, the zero-control infinitesimal generator, **A**_0_, is constructed according to:

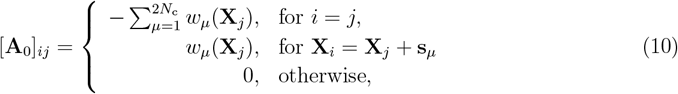

and the feedback infinitesimal generator, **Bu**^*𝒞*^, of the controller is constructed according to

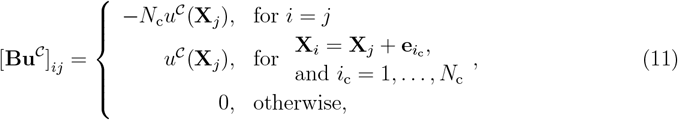

where *u*^*𝒞*^(**X**_*j*_) is the specification of the controller, *𝒞*, in terms of the current instantaneous state, or its partial observations.

For a given controller, the equilibrium distribution of the system (**P**^*∗*^) can be found by solving Eq. 9 and is given by:

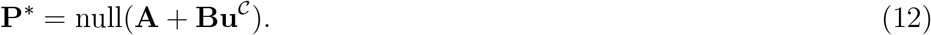

In principle, the master equation in Eq. 9 could contain an uncountably infinite number of states, and therefore the exact solution as well as the null vector in Eq. 12 may not be computable exactly. To address this issue, we first truncate the system at a finite number for each species and then apply a reflecting boundary condition, resulting in a finite dimensional master equation.

### 2.3. Quantification and optimization of control performance

In the SIMO control of stochastic processes, a single input is applied simultaneously to all systems at once, and therefore a control signal that acts beneficially on one cell may destabilize other cells. An effective controller must strike a balance among the desired behaviors of all cells in the system. To quantify overall performance success, we define the *steady state performance error, J*, as the expected steady state Euclidean distance of the process from the specified target state, **T**:

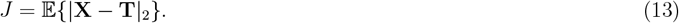

The squared score is easily calculated by applying a linear operator to the steady state probability distribution **P**^*∗*^ (Eq. 12) as follows:

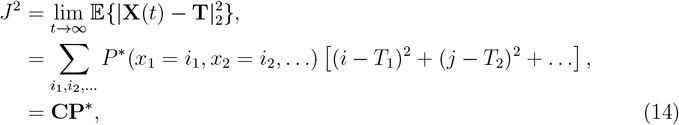

where **C** is simply a vector that contains the squared Euclidian distance of each state from the specified target **T**, i.e., 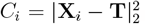. As a result of this calculation, *J* is a non-negative scalar that is zero only if **P**^*∗*^ is a delta distribution located exactly at the target vector **T**.

We consider two controller designs: the fully aware controller (**u**^*FAC*^) that bases its control signal on simultaneous protein count observations from both cells, and the partially aware controller (**u**^*PAC*^) that relies only on observations from a single cell:

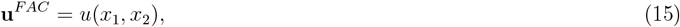

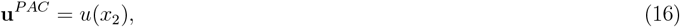

where *x*_1_ and *x*_2_ are discrete integers greater than or equal to zero that represent the instantaneous number of proteins in cell one and cell two. Despite their differences in their observation data, both the FAC and the simpler PAC are optimized to achieve the same goal, namely to drive both cells to their respective set points.

Optimization of **u**^*F AC*^ and **u**^*PAC*^ were performed using a gradient descent method to minimize *J*. Since the square root is a monotonically increasing function, minimizing*J* ^2^ results in the same control as would minimizing *J* directly. Therefore, we calculate the negative gradient 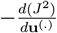 and adjust parameters a small step *d***u**^(.)^ in that direction. For example, the calculation of the gradient for the FAC controller is given by:

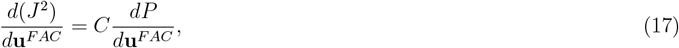

Where 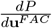 can be solved using general minimized residual calculation of

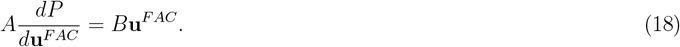

In [23], controllers were optimized to minimize *J* ^2^ for a single target state **T** = [10, 30], but in this work they are extend to different arbitrary target points.

### 2.4. Scaling for system granularity

In realistic applications, models are never exact, but are often chosen as simplifications of known processes. For example, when analyzing discrete stochastic chemical kinetics, it is common to project the CME onto lower-dimensional spaces using finite state projection [26], time scale separations [27], Krylov subspaces [28], principle orthogonal decompositions [29], or other coarse meshes [30, 31]. Similarly, measurements are also always inexact and in many cases may only provide information about relative changes – for example, although fluorescent proteins usually cannot be counted exactly, one may reasonably assume that a cell’s total fluorescence intensity varies linearly with the fluorescent protein concentration. To explore how mismatches in the assumed system scale (e.g., arising from model approximations or relative measurements) affect the controllability of the cellular process, we define a granularity parameter (*α* = *M*^′^*/M*) that linearly scales each species’ population to increase (*α >* 1) or decrease (*α <* 1), while maintaining the dynamics and general behavior of the model. To apply this granularity parameter, we assume that each propensity function, *w*_*μ*_ (Eqns. 4) is rescaled to a different level of discreteness by substituting

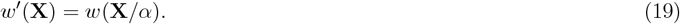

For example, the production and degradation of protein in cell one would become:

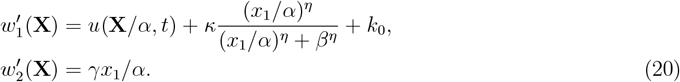

We note that in order to reuse a control law that has been defined for one level granularity and apply it to a system at another level of granularity, the inputs to the controller must also be scaled by 1*/α* before computing the control level assigned to the current state, e.g., *u*^*𝒞*^ = *u*^*𝒞*^(**X***/α*). Because identification of the original control formulation, **u**^*F AC*^(*x*_1_, *x*_2_) and **u**^*PAC*^(*x*_1_) only considered integer values for (*x*_1_, *x*_2_), control signal values at fractional state values after rescaling (*x*_1_*/α, x*_2_*/α*) are calculated using 2D cubic interpolation from the control values at the nearest integer state values. Finally, to provide a consistent metric for relative scoring, the definition of the performance score is also adjusted according to scale magnitudes. For example, in the two cell system the new steady state performance error would become:

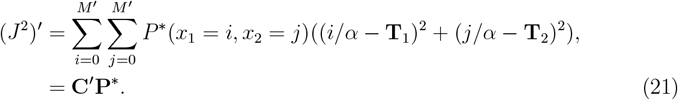

We reiterate that the system parameters and control law were defined and fixed using the the base granularity (*α* = 1), and to simulate a practical application where scales may be unknown or variable, these are not recomputed or refit upon changing the system granularity.

### 2.5. Observation and actuation time delays

Delays are inherent to any realistic control system, and in this case delays would be expected to arise due to the time needed for various biochemical reactions such as the formation of complete polymerases, activation of promoters, transcription and transport of mature mRNA, and the translation and maturation of protein [32]. Additional delays would also arise from data analysis, decision making, and actuation dynamics. To investigate the effects of observation or actuation time delays on control performance, we devised a simple time-delay stochastic simulation algorithm. This algorithm records the state history after each stochastic reaction, enabling reconstruction of the population history of the species and the time delayed control input propensity. Using this information, the time-delayed control signal at time *t* can be specified as:

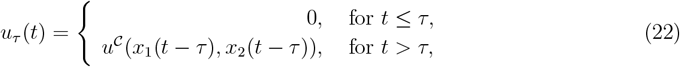

where *τ* is the time delay between observation and actuation, *u*^*𝒞*^ is the previously optimized control law (e.g., **u**^FAC^ or **u**^PAC^), and *x* gathered from the state history. We note that the time delay stochastic process was only simulated using the SSA because to our knowledge an appropriate direct FSP/CME integration procedure has not yet been developed. The Extrande method [33] was used to update the control input at an average frequency of 50 updates per minute, far exceeding the dynamics of the system.

### 2.6. Tracking Time-Varying Trajectories

Because optimizing the control law for a static set point as described in Section 2.3 requires differentiation of the CME (Eq. 9) with respect to the control signal at each state, this calculation is approaching the limits of current feasibility. Extending these calculations to optimize controllers for a dynamically moving set point is much harder and would likely require intractable numerical simulation or development of new mathematical approaches that are beyond the scope of the current study. To circumvent this challenge, we instead propose a simple alternative in which the controller sweep though a piecewise constant set of controllers each designed for a specific static target point along the desired trajectory.

Our goal is to control the system to follow a specific target trajectory, **T**(*t*). We choose a discrete set of *K* target points along this trajectory:

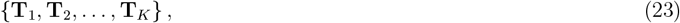

and for each individual target **T**_*i*_, we optimize to find a corresponding controller, **u**_*i*_ = **u**(**T**_*i*_). Finally, to implement the control at any given time, *t*, we find the index of the nearest precomputed target state:

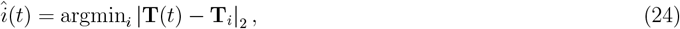

and assign that controller. Under this piecewise constant controller law, the full time-varying CME becomes

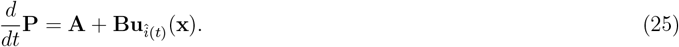

### 2.7. Code and data availability

All codes and data needed to reproduce figures is provided at https://github.com/MunskyGroup/May_2023_Exploiting_Intrinsic_Noise. A frozen version of this repository will be provided via Zenodo upon acceptance of the manuscript.

## 3. Results

Using the analyses described in the Methods section, we focus on a single-input-multiple-output (SIMO) controller where *all cells receive the same input* at every instant of time. For this, the control signal depends upon the observed state, e.g., *u*(*t*) = **u**(**x**) as illustrated in Fig. 1(I,III). These controllers have been optimized to break symmetry so that multiple cells can be controlled to different target pstates as illustrated in Fig. 1(II,IV), and we quantify performance according to the RMSE error introduced in Eq. 14. To explore how such controllers may perform in realistic settings (e.g., where the models are approximate or the parameter are unknown or extrinsically variable), we now fix the parameters of those controllers and explore performance robustness to different types of model uncertainties, including parameter errors or variations (Section 3.1), incorrect assumptions on system scales (Section 3.2) and time delays (Section 3.3). Finally, in Section 3.4, we extend the control analysis to consider the performance of the controller for tracking variable reference signals with different frequencies and phases.

### 3.1. Stochastic SIMO Optogenetic control can remain effective despite small parameter errors or extrinsic uncertainties

Real stochastic processes always have unknown or uncertain mechanisms and parameter values. Although model structures and parameter estimates can often be obtained through fitting to training data, these estimates will never be perfect due to unavoidable measurement errors or limited data sets. Even with plentiful and precise training data, model and parameter uncertainties are inevitable due to the fact that even genetically identical single-cells exhibit heterogeneity in parameters due to extrinsic variations. To asses the control performance under such parameter errors, we performed sensitivity analysis on each model parameter for each cell, then on both cells simultaneously. For each parameter, **u**^FAC^ (solid lines) and **u**^PAC^(dashed lines) control performance was examined across a parameter perturbation range from one-tenth to ten-fold of the original values. Figure 2A shows the results for these parameter sweeps when a single parameter in Cell 1 is varied (all other parameters are fixed), and Figure. 2B shows the control performance when a single parameter in Cell 2 is varied. Finally, 2C shows the performance change when a single parameter is changed simultaneously in both Cell 1 and Cell 2 at the same time.

**Figure 2.**
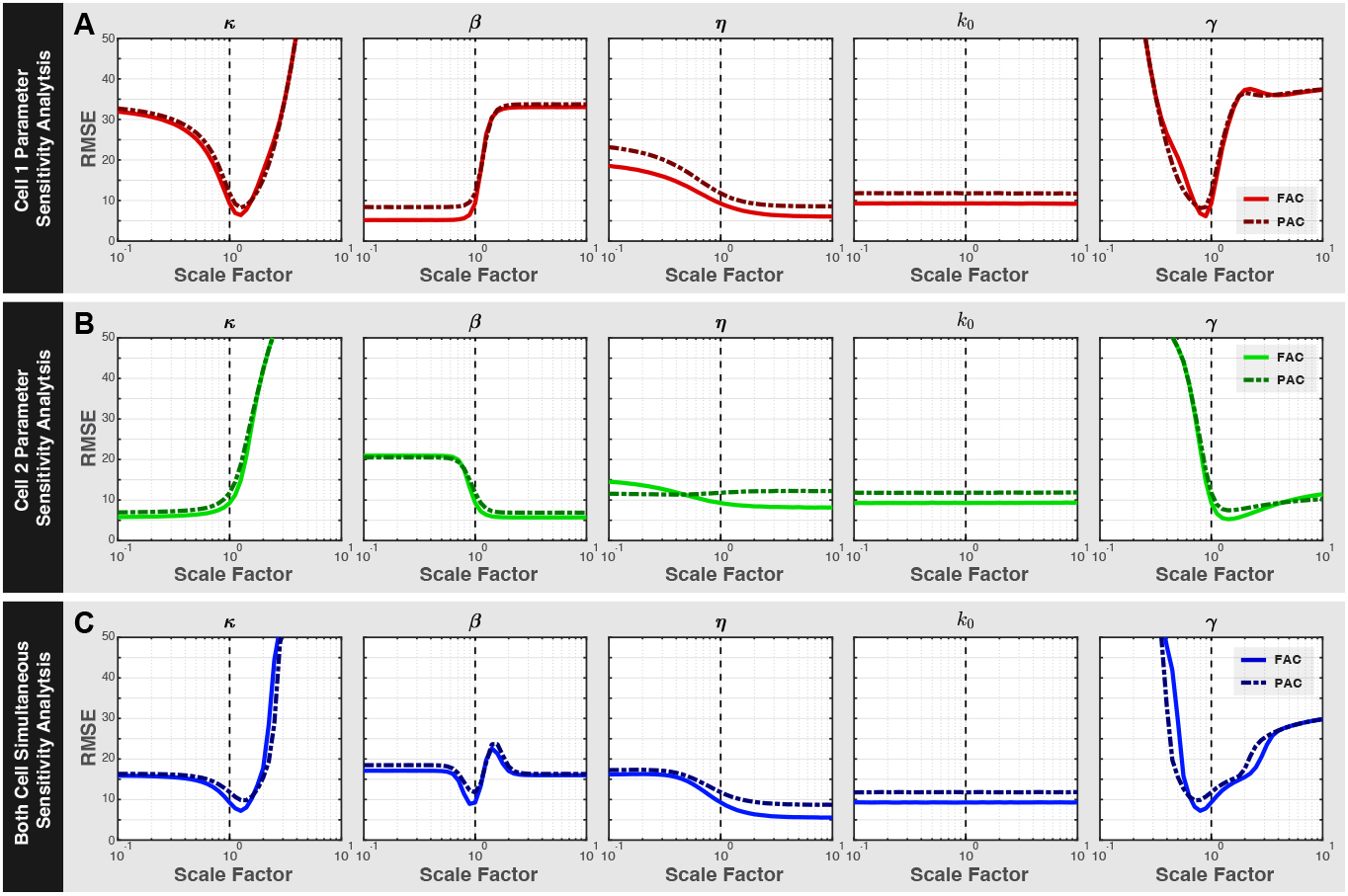
Parameter sweeps using the FAC and PAC show a broad range of control performance in cell 1 (**A**), in cell 2 (**B**), and in both cells (**C**). Columns show each parameter in the model, rows show the cell which has its parameter perturbed.

Modifications of parameters were found to produce a broad range of effects. For example, increasing *β* in Cell 1 quickly worsens performance while increasing *β* in Cell 2 improves performance (compare second column in Figure. 2A and 2B). In some cases, the effects on performance are not monotonic; for example, increasing *κ* in Cell 1 (Figure. 2A, leftmost column) would be highly advantageous up to a limit after which the control performance degrades rapidly. In other cases (such as for the promoter leakage rate, *k*_0_), the effect of parameter perturbations on performance is insignificant even for relatively large (*s*=10) perturbations, suggesting that this parameter is not important to control performance. Figure 2C shows that even when parameters of both Cell 1 and Cell 2 are jointly changed, these changes could also improve or detract from control performance. In particular, the analysis shows that control performance could be improved by modifying the system design either to increase the auto-regulation promoter strength (*κ*) or its cooperativity (*η*) or to decrease the promoter binding constant (*β*) or the protein degradation rate (*γ*). We note that co-optimization of both the system and the controller law would allow for further improvements to the performance.

### 3.2. Controllers trained using one assumed level of granularity can remain effective at other levels of granularity

We next explored how the granularity of the system would affect the differential control of the multi-cell system. Specifically, we define the level of granularity using the parameter *α* as described in Section 2.4 and which affects the propensity functions and scoring according to Eqns. 19-21. Effectively, larger *α* corresponds to systems where larger numbers of individual molecules are needed to achieve the same concentration (e.g., larger volumes), while smaller *α* corresponds to situations where smaller population sizes can achieve that concentration (e.g., smaller volumes). We previously optimized the controllers **u**^FAC^ and **u**^PAC^ based on a the default assumption that *α* = 1, and we wished to know what would be the consequences if this same controller were to be applied to a system that has a different granularity.

To explore this tradeoff, control performance scores for the **u**^FAC^ and **u**^PAC^ controllers were calculated at different levels of granularity (*α*) between 0.2 and 2.0. Figure 3 (A -F) shows the joint probability distributions (left plots) and marginal probability distributions (right plots) of the system at a low granularity (*α* = 0.2), the original granularity (*α* = 1.0), and at an increased granularity (*α* = 2.0). Figure 3G shows the trend of the performance versus alpha for both the FAC (solid cyan line) and the PAC (solid magenta line) controller. This improvement in performance appears to approach a small value as the granularity goes to infinity, but since the size of the FSP increases with the square of the system size, systems much larger than *α*=2 (where 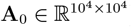) become more difficult to calculate using master equation techniques. To bypass this limit in the FSP, sixteen SSA simulations were used to sample the CME of a system with a much larger volume of *α*=100. Each SSA was run for 5 *×* 10^7^ minutes and only the last 4 *×* 10^7^ minutes were sampled to estimate the stationary distribution and calculate the performance score. The performance score estimates of this high granularity SSA using the FAC and PAC were 2.03 and 5.24 respectively, which are plotted as dashed lines in Figure. 3G. Although it is unclear if further performance improvements could be obtained with further increases to the system scale, for all cases considered so far, we found that both controllers monotonically improved with increased *α* and that **u**^FAC^ always outperforms the **u**^PAC^.

**Figure 3.**
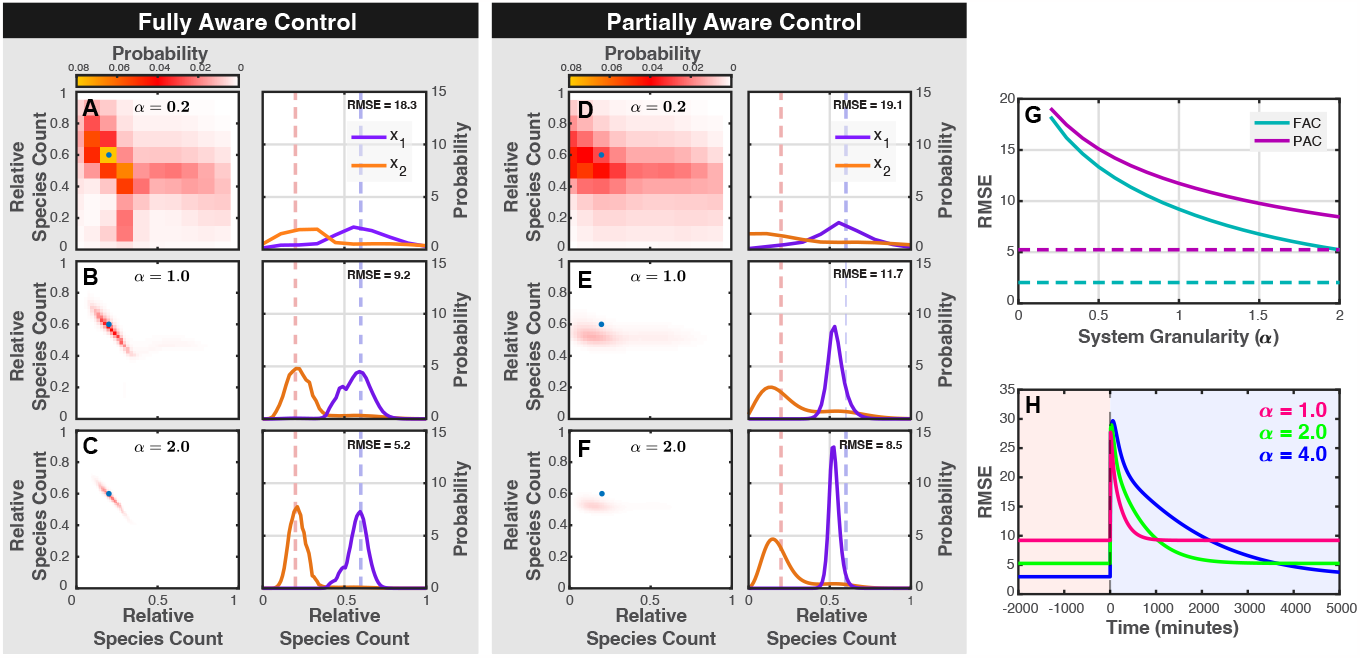
Systems with increased granularity are less noisy and have better control performance. (A-C) Joint (left) and marginal (right) distributions for the FAC shows increased control performance and tighter distributions as *α* increases from 0.2 (top row) to 2.0 (bottom row). (D-F) Joint (left) and marginal (left) distributions using the PAC controller at different levels of granularity.

The effect that granularity has on the control performance depends on two competing phenomena: First, because the controller was optimized for one level of granularity (*α* = 1), one might expect that the controller would become worse if the granularity were incorrect. However, at large granularity, we find that the opposite is true -the control performance actually improves when applied to an incorrect model. The reason for this surprising result is that relative amplitude of stochastic fluctuations (i.e., the standard deviation divided by the mean) in a chemical process decreases with the inverse square root of the process scale [34]. In other words, the process becomes more predictable and therefore more controllable. At the extreme as the system size increases, the dynamics converge towards a deterministic process, except for certain exceptional initial conditions lying on manifolds that would separate different steady state behaviors [35].

However, this improvement in the steady state performance does not come without a cost. Although higher granularity reduces noise and makes it easier to maintain desired states once they have been achieved, noise is necessary to break symmetry between the two cells’ dynamics in order to achieve those states in the first place. This tradeoff is illustrated in Fig. 3(H), which plots the control performance over time after changing the control goal to exchange the low- and high-target cells with one another. From the figure, we can see although steady state control performance improves for higher values of *α*, the time taken to reach that steady state performance increases with *α*, suggesting that larger-volume system may become more susceptible to times delays or less able to track variable reference trajectories.

### 3.3 Heterogeneous control can remain effective despite moderate time delays

In general, feedback control can only be effective if one can quickly make measurements, compute adjustments to the control signal, and implement the needed changes within an appropriate amount of time relative to the characteristic timescale of the system. As the time required for any of these steps increases, control performance will be degraded, perhaps even leading to large fluctuations or instability. To explore how time delays affect the noise-enhanced controllers **u**^FAC^ and **u**^PAC^, we generated large sets of time-delayed stochastic simulations (see Section 2.5) for different lengths of the time delay. Each SSA was sub-sampled for 1000 times over 10000 minutes of simulation time after a burnin period of 10000 minutes.

Figure 4 shows the joint distributions (left) and marginal distributions (right) at varying levels of time-delay, with panels A-C showing results for the FAC controller and panels D-F showing results for the PAC controller. Figure 4G summarizes these results by plotting the score of both controllers versus the time delay. From the figures, it is clear that performance is rapidly degraded as the delay approaches and then exceeds the characteristic time (*τ*_*𝒞*_ = 1*/γ* = 49 min) set by the degradation rate of the process (yellow dashed line). At very small time delays (*τ <* 3.4 min), the FAC outperforms the PAC, but at moderate time delays (*τ >* 3.4 min), the PAC outperforms the FAC. At high time delays, both controllers lose their asymmetry, resulting in substantially worse performances (*RMSE* = 32 and 23) and even perform worse than under a simple constant control input without feedback (depicted by a horizontal red line in Fig. 4G). As discussed in the previous section, increasing the granularity of the system *α* reduces the randomness to improve the steady state performance but at the cost of slowing down the controller response. To explore the joint effects of time delays and granularity, Fig, 5J shows the FAC steady state control performance as a function of the time delay and system granularity (*τ, α*) using thirty-two stochastic simulations simulated to steady state at each combination. Control performance errors measured by RMSE improved as *α* increased when the delays were small (*τ* = 1 min and *τ* = 10 min) but not when *τ* = 100 min.

**Figure 4.**
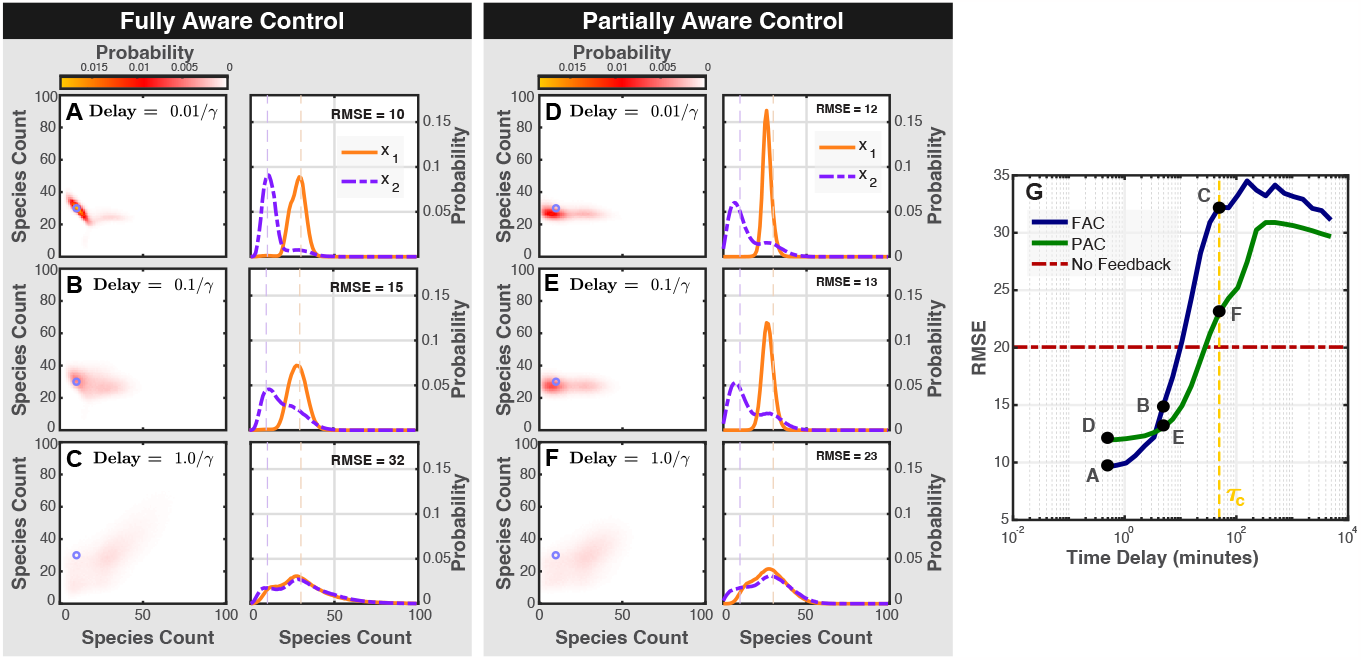
Effects of time delay on control performance. (A-C) Joint (left, color scale shown at top) and marginal distributions (right) of the controlled system at different levels of time delay using the FAC controller. The target state, **T** is denoted by a small circles on the left panels and dashed lines on the right panels. The steady state RMSE is shown in each case. (D-F) Same as (A-C) but for the PAC controller. (G) RMSE control performance versus time delay for both FAC (blue) and PAC (green). Letters A-F correspond to panels A-F. Dashed red line corresponds to optimal performance with no feedback (i.e., constant input). Dashed yellow line corresponds to characteristic system time, *τ*_*𝒞*_ = 1*/γ*.

Recalling that under the system granularity rescaling (Eq. 20), as *α* changes, the effective degradation rate scales according to *γ*^′^ = *γ/α*. Therefore, the critical limit for the delays should also change with the system scale according to *τ*_*𝒞*_ = *α/γ*. Figure5(J) depicts this characteristic line and shows that as the time delay approaches and then exceeds this level, the steady state performance becomes dramatically worse.

Finally, to understand the model of failure at these longer delays, it is interesting to examine trajectories induced by the controller just below this characteristic delay. For parameter set denoted by the red star in Figure. 5(J), Figure. 5(K) shows the controlled response after a long burn in period to achieve steady state. In this case, the application of feedback after a delay leads to a strong oscillatory behavior and worse performance than that achieved without any control at all. This observation further stresses the importance of considering time delays when designing such controllers.

**Figure 5.**
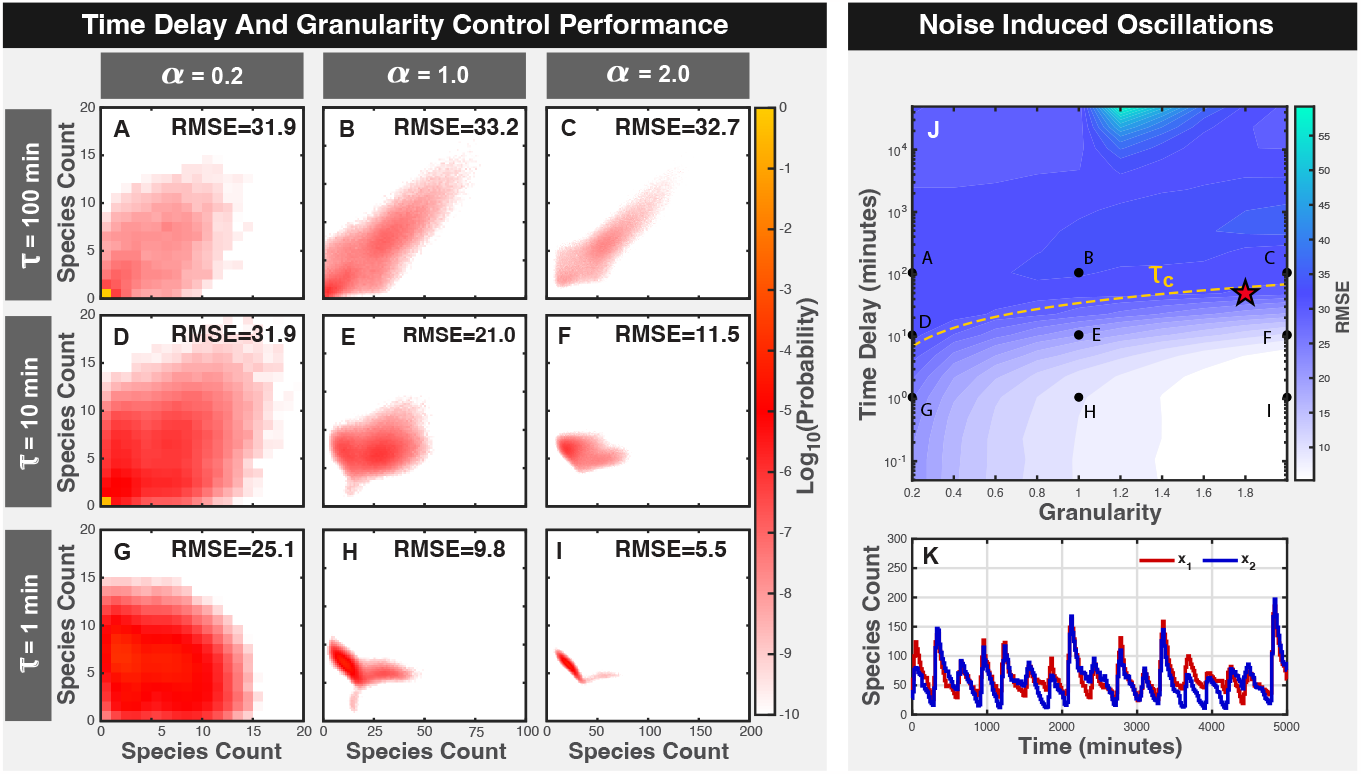
Joint effects of granularity and time delay on control performance. (A-I) Joint probability distributions at different combinations of *τ* (rows) and *α* (columns). Overall RMSE shown at top. (J) Heat-map of control performance versus (*τ, α*). Points A-I correspond to panels A-I. Dashed yellow line shows characteristic time *τ*_*𝒞*_ = *α/γ/*. (K) Controlled system trajectory for *α* = 4.0 and *τ* = min (denoted by red star in panel J).

### 3.4. With noise-induced control, a single input can drive multiple cells to follow different temporal trajectories

We next optimized the FAC controller and calculated its performance for the two-cell system over a two dimensional domain of discrete target points **T**_*i*_ = [*T*_*i*1_, *T*_*i*2_] between between five and forty-five. For example, Fig. 6 (A) shows the optimized FAC control input of the system when the target is **T** = [20, 25], and Fig. 6 (B) shows the corresponding steady state probability distribution of the controlled system. Figure 6 (C and D) show the the same thing but for a different target state of **T** = [10, 25]. Figure 6(E) shows the overall steady state FAC control performance over the entire domain of static set points, and illustrates that some regions are easier to attain than others. For example, the top-left and bottom-right edge regions (i.e., where one species is well above its high equilibrium point and the other is well below its low equilibrium point) are the most difficult to control, while other target points near the two equilibrium points are more easily obtained.

**Figure 6.**
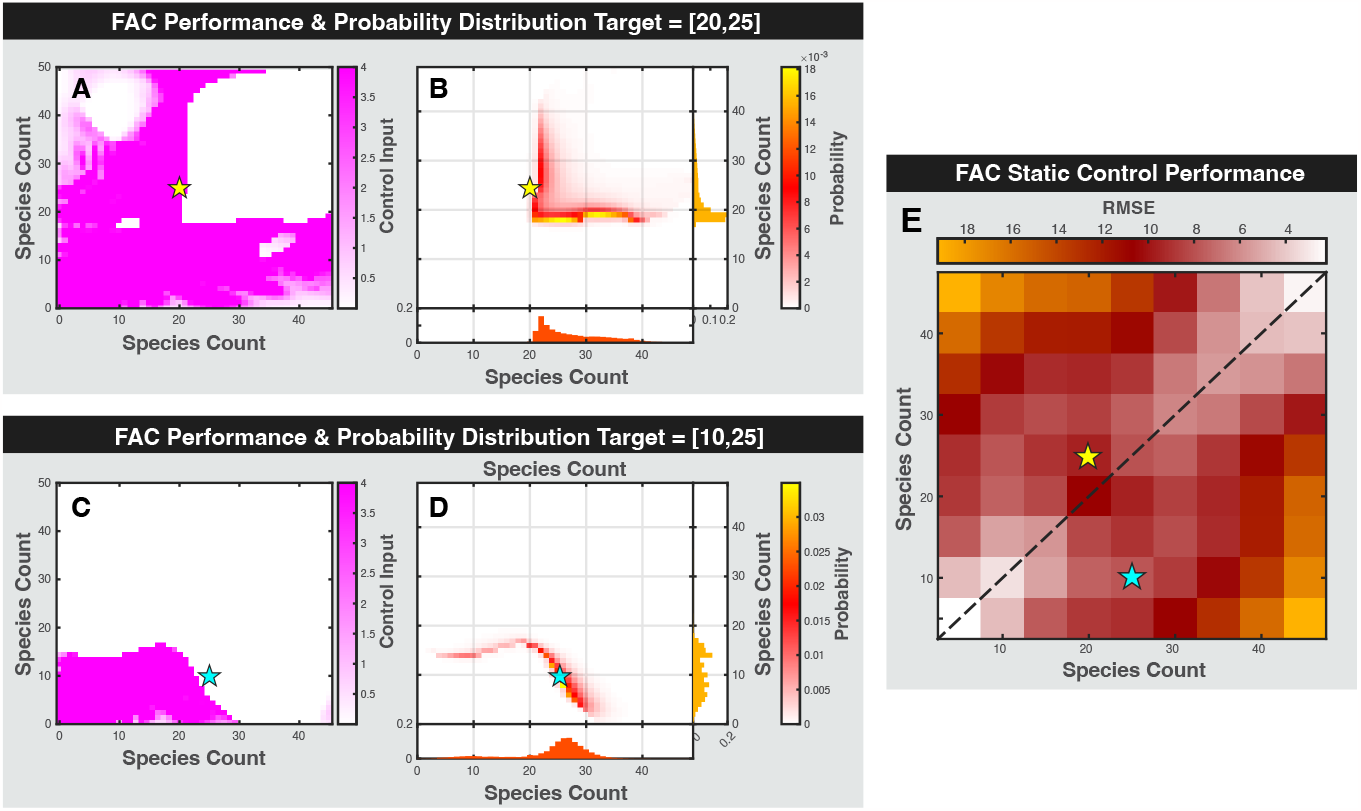
FAC control laws and performance for different target points. (A) Optimized control input for target **T** = [20, 25], with target point denoted by star. (B) Corresponding steady state response distribution. Marginal distributions for *x*_1_ and *x*_2_ on right and below. (C,D) Same as (A,B) but for **T** = [10, 25]. (E) FAC control performance (RMSE) versus targets **T** = [*T*_1_, *T*_2_]. Stars correspond to target points in panels A-D. Dashed diagonal shows line of symmetry.

We developed a method (Section 2.6) to control the system dynamics to follow a predefined path **T**(*t*) by alternating between 32 different pre-computed controllers along each path. We considered three representative pathways, including an in-sync reference point (Fig. 7B1) where

**Figure 7.**
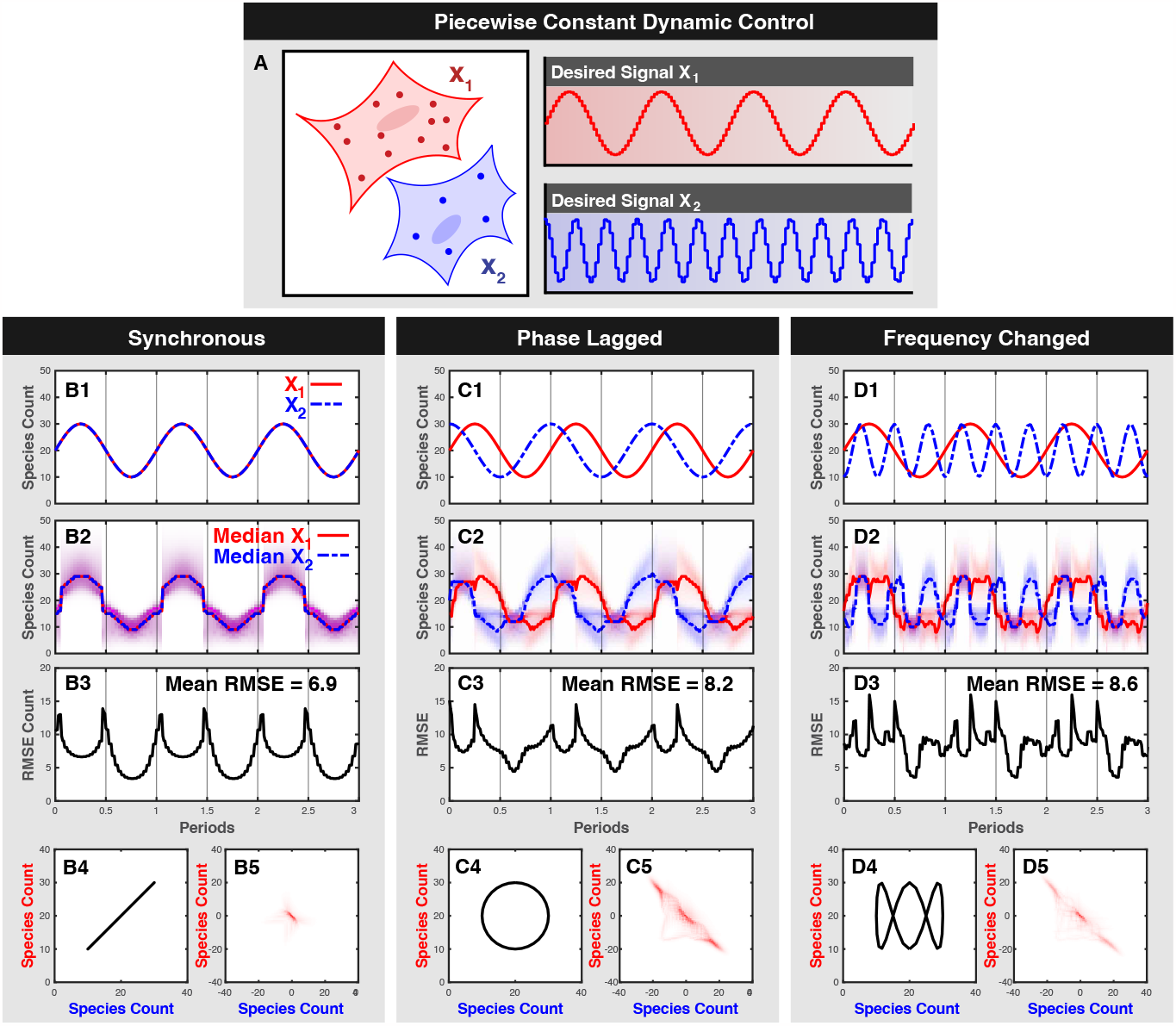
Tracking time-varying reference signal. (A) Schematic of SIMO control to drive two cells to follow different trajectories. (B1) Reference signal for *x*_1_ (red) and *x*_2_ (blue). (B2) Controlled response. Distributions shown in shading. Median shown in lines. Three periods are shown after decay of transient dynamics. (B3) RMSE performance over time. (B4) Phase space of reference signal. (B5) Time-averaged distribution of tracking error. (C1-C5) Same as (B1-B5) but for phase-lagged reference signal. (D1-D5) Same as (B1-B5) but for reference signal with two different frequencies and phases.

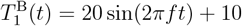, and 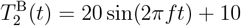;

a phase lagged reference point (Fig. 7C1) where

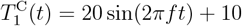, and 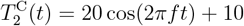;

and a frequency separated reference point (Fig. 7D1) where

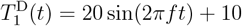, and 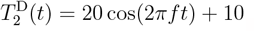.

For each reference signal, a slow driving frequency was applied at *f* = 10^*−*4^ cycles per minute.

All FSP simulations were calculated under the time varying control law, and Figs. 7(B2, C2, and D2) show the corresponding response distributions (shading) and median responses (lines) for two cells *x*_1_ and *x*_2_ in red and blue, respectively. Regions with purple shading depict the exact overlap of red and blue, when both cells have the same distribution of response. From Figs. 7(B2, C2, and D2), we observe that the SIMO control system can effectively drive the system to follow all three input trajectories, including when the two reference trajectories have different phases and frequencies (panel D2). To quantify the overall performance, Figs. 7(B3, C3, and D3) show the root mean squared error as a function of time, and Figs. 7(B4, C4, and D4) show the time-averaged error distribution for the system response relative to the time varying target. From these figures, we observe that all trajectories result in short RMSE spikes during short transient periods when the controller passes through the regions of poor control (i.e., through regions found in Fig. 6E to have high RMSE errors). Overall, the simulations show that the synchronous control performed best with an average RSME of 6.9. When the system is driven with a phase-lag, the average RSME of the score increases to 8.2, and when driven at a different frequency, the average RMSE goes up to 8.6.

In general, a given control system cannot effectively track a reference signal that changes faster that the system’s natural time scale. Since it was shown that increasing granularity (*α*) improves steady state control performance (Fig. 4G) but also lengthens this time scale (Fig. 4H), one should expect that these competing effects of granularity would also affect the types and speeds of signals to which the system can respond. To examine how driving frequency and system scale affects control performance, 64 SSA trajectories of the phase-separated dynamic controller were simulated over a two dimensional domain of points (*α, f*) for 32 cycles after reaching steady state. Figure 8(A) shows the desired reference signal for the system, and Fig. 8(B) shows the mean of the system response when the frequency is *f* = 10^*−*4^ cycles/min and the granularity is *α* = 4.0. Figure 8(C) shows the corresponding control performance over normalized periodic time as *α* is held at 4.0 and *f* is increased from *f* = 10^*−*4^ cycles/min to *f* = 10^*−*2^ cycles/min and *f* = 10^*−*0^ cycles/min leading to average control performances of 6.2, 16.1, and 15.0 respectively. Figure 8D,E extends this analysis to plot the average control performance over different combinations of *α* and *f*. The characteristic frequency (given by the predicted 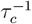 over a range of *α*) is plotted as the yellow dashed line. In particular, the heat map of control performance over (*f, α*) shows that the worst performance occurs at moderate *f* near *f* = 10^*−*2^ cycles/min and that control performance is best when granularity is high and frequency is low 8(E). More specifically, we find that strong performance was attainable only if the driving frequency was kept lower than a characteristic frequency *f*_*𝒞*_ *≡* 1*/τ*_*𝒞*_ = *γ/α*, which is denoted by the dashed line in Fig. 8E.

**Figure 8.**
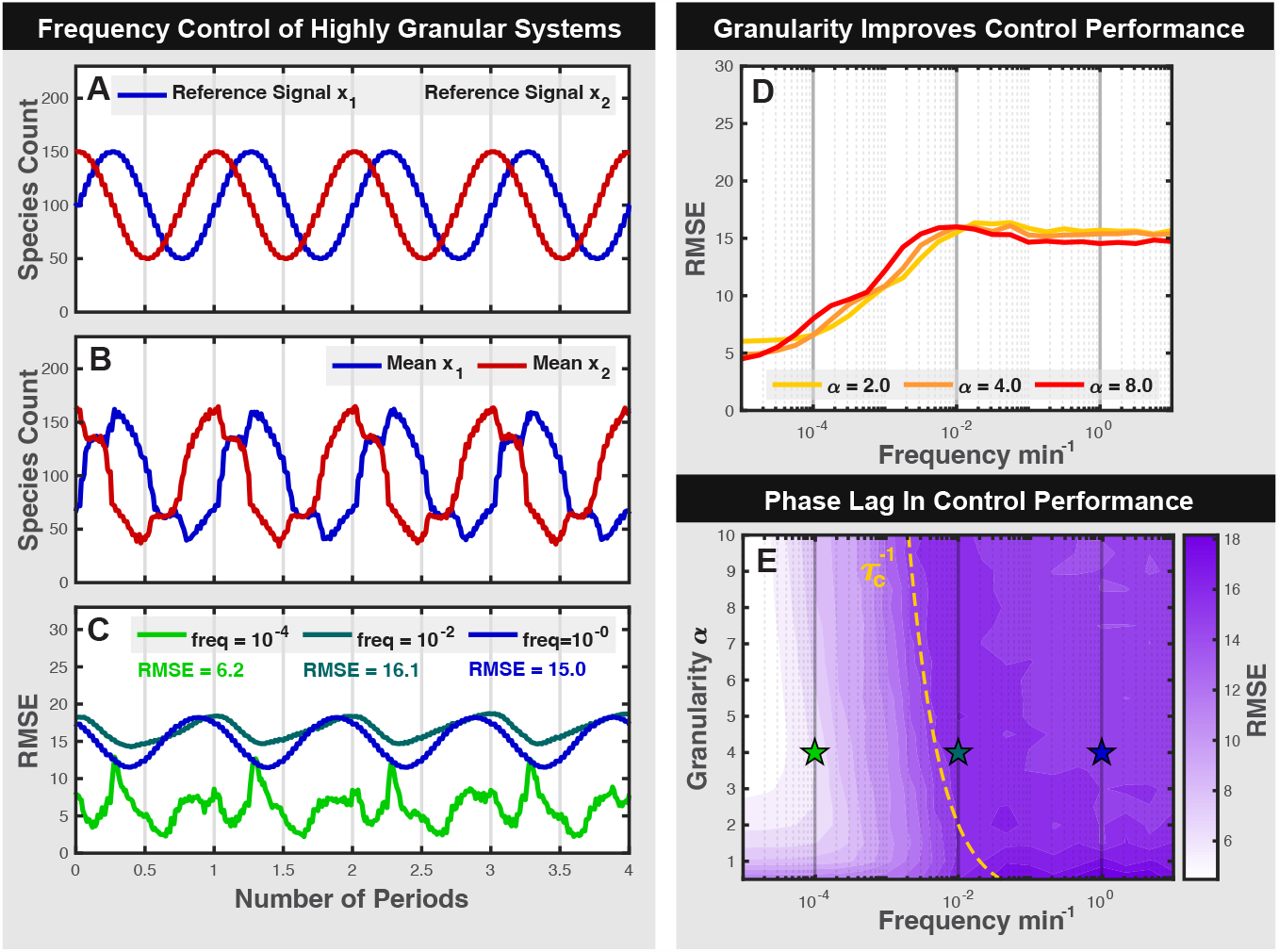
Control performance analyzed over a domain of *f, α* pairs show worst control performance at moderate frequencies near 1*e −* 2 due to phase lag. Tracking reference signals when *α* = 5 span a range between 50 and 150 species (A). Stochastic simulations driven using a phased lagged controller at *α* = 5 and low frequency show tighter control compared to *α* = 1 (B). Systems driven at moderate frequency show worse control performance than high frequency or low frequency (C). Control performance only of phase-lagged system only improves with increasing *α* and low *f* .

## 4. Conclusion

Noise, whether it arises from inherently stochastic processes or from unknown or unmodeled interactions, can play a critical role in the performance of feedback control. For the field of synthetic biology, this noise has typically been avoided and genetic systems have primarily been engineered to be as robust as possible to these uncertain fluctuations. In contrast, many natural cellular processes exist and thrive in settings where single-molecule events such as gene activation leads to large relative fluctuations, and where response heterogeneity is unavoidable. Key results from [23] showed that certain controllers could exploit this noise to achieve objectives that would not be possible in a deterministic setting. With such controllers, a single regulatory signal, such as an optogenetic input, could drive two or more two genetically identical cells to different, arbiltrarilly chosen fates using just a single input signal and irrespective of the cells’ initial conditions.

The effectiveness of any model-based controller depends upon the accuracy of the model with which that controller has been optimized, and as the real system deviates from its idealized model, the control performance will naturally be affected. In this work, we explore the effects of several such deviations, including uncertainties or errors in parameters, mismatches in assumptions for systems scale, and times delays.

Regarding parameter values, our perturbation analyses (Fig. 2) showed that control performance is strongly affected by the system parameters. We found that whether parameters were incorrect in all cells (e.g., due to systematic errors in the model) or for just one cell at a time (e.g., due to extrinsic noise in the cells themselves) affected the control performance in different way. In most cases, small changes could be tolerated, whereas large changes, especially to certain key parameters, could be catastrophic. Moreover, we found that there can be room to improve control performance by adjusting system parameters, suggesting that joint optimization of the controller with the system itself could lead to even stronger performance. Armed with such insight into which parameters are the most sensitive and which can safely be ignored, one could in principle focus measurement efforts to more precisely quantify the critical parameters and focus design efforts to reduce variability in key aspects of modular parts.

The size of the system also plays an important role in its control performance. For a fixed concentration, as the volume of a chemical reaction system increases, it becomes less noisy, and its dynamics approach that of deterministic process. At this limit, symmetry can no longer be broken, and feedback control cannot independently drive different cells to different fates. On the other hand, deleterious fluctuations also become smaller, so larger systems can more easily maintained maintain their desired phenotypes. Overall, we have shown (Fig. 3A-G) that the removal of noise through system granularity led to better steady state control performance, but such systems were found to take a much longer time to achieve steady state (Fig. 3H). Interestingly, this result could have implications on the malleability of cells at different stages of their growth cycle, where differentiation of smaller cells (e.g., those immediately after division) may be more susceptible to control signals, while larger cells (e.g., mature cells that have already established their phenotypes) may be relatively impervious to external signals.

Although our objective was to determine how well control performance would be maintained under different system sizes, we were surprised to find that controllers designed at one level of granularity (e.g., *α* =1) worked surprisingly well to control systems at much larger granularities. From a practical perspective, the ability to analyze a model at one system scale and then effectively apply it to another could be highly beneficial. Since the computation time of the FSP solution to the CME grows with the square of the number of states, this can cause an explosion in computational requirements for large systems. Our results suggest a promising alternative in which one could learn or optimize a controller using FSP analyses for a computationally feasible number of states and later apply them to larger systems that cannot be solved using current techniques.

Time delay analysis (Fig. 4) showed that increasing time delay decreased control performance. However, we also found that not every controller was equally affected by time delays. In particular, we found that at intermediate and larger time delays, a partially aware controller that has less information can outperform a fully aware controller (Fig. 4G). We believe this is happening because a controller with more information can afford to be more aggressive to implement its control, and time delay can cause this aggression to backfire.

The control of cells to two slowly changing dynamic reference signals using a single global input by the use of a noise-exploiting controller showed good control performance for a variety of signals. These analysis could be extended to include faster frequency by numerical calculation or by alternative error-probability adjustments.

## Notes

### Competing Interest Statement

The authors have declared no competing interest.

